# Regulation of Fungal Morphology, Conidiogenesis and Violet Pigment Synthesis by the Betalain Biosynthesis Pathway Genes in *Aspergillus sydowii* H-1

**DOI:** 10.1101/2024.12.13.628389

**Authors:** Yulu Ran, Yu Cao, Yihan Guo, Jie Zeng, Jiale Wang, Dongyou Xiang, Hui Xu, Yi Cao

## Abstract

The biosynthesis of antioxidant pigments, namely, betalains, has predominantly been found in Caryophyllales. The potential betalains biosynthesis was firstly explored in *Aspergillus sydowii* H-1 under controlled culture conditions. This study identified, knocked out, and overexpressed genes involved in the betanin biosynthesis and assessed the activities of tyrosinase, 4’5-DOPA dioxygenase and LigB. The results indicated these betanin biosynthesis pathway was crucial for colony morphology, conidiogenesis, stress response, and violet pigment synthesis. Moreover, AsDODA and AsLigB catalyzed the conversion of L-DOPA into 4’5-seco-DOPA, a key intermediate in the formation of betalamic acid in vitro. Additionally, transcription factors such as AsbHLH, AsMYB1R, and AsWD40 positively regulated the expression of betalain biosynthesis genes. This research provides new insights into the evolutionary origins of betalain-producing species, expanding the scope of betalain biosynthesis to include *Aspergillus* species.

**Importance:** To date, betalains were restricted to plants of the order Caryophyllales, fungi of Basidiomycota and several types of bacteria. This study is the first to demonstrate the potential of Ascomycetes *A. sydowii* H-1 to synthesize betalains under controlled culture conditions, providing a detailed genetic and biological characterization of the associated genes and metabolic pathways. This finding demonstrates that betalain biosynthesis can be expanded to other *Aspergillus*.

## Introduction

Secondary metabolites are a variety of low-molecular-mass compounds synthesized by fungi, essential for various cellular processes including transcription, development, and intercellular communication[1]. *Aspergillus sydowii* is a saprotrophic, ubiquitous, and halophilic fungus prevalent in marine ecosystems. Remarkably, *A. sydowi* synthesis a wide array of metabolites, including alkaloids, xanthones, terpenes, anthraquinones, sterols, and so on, each exhibiting various bioactivities and crucial applications[2]. In 2016, we successfully isolated and identified a fungus strain *A. sydowii* H-1 from Chengdu, China, which produced a violet pigment during fermentation, exhibiting antibacterial, antioxidant, and antitumor activities[3].

Betalains are pigments(a kind of secondary metabolites) categorized into two groups, the violet betacyanins and the yellow betaxanthins, which in addition present green fluorescence[4]. Betalain biosynthesis consists of three key steps. First, tyrosine is converted to L-DOPA (L-3,4-dihydroxyphenylalanine), catalyzed by CYP76AD or tyrosine hydroxylase. Next, L-DOPA 4,5-dioxygenase (DODA), a member of the LigB gene family, cleaves L-DOPA to form the intermediate 4,5-seco-DOPA via a ring-opening oxidation reaction. This is followed by the spontaneous intramolecular condensation of 4,5-seco-DOPA to produce betalamic acid. Simultaneously, L-DOPA may be oxidized to dopaquinone, which cyclizes to form cyclo-DOPA, a process catalyzed by CYP76AD. Finally, betalamic acid conjugates with the imino group of cyclo-DOPA, leading to the formation of violet betacyanins. Alternatively, betalamic acid may condense with the imino or amino group of amino acids, resulting in the formation of yellow betaxanthins[5]. Betalains play important roles in attracting animal pollinators and dispersers, contributing to photoprotection, and enhancing tolerance to drought and salinity stress due to their high antioxidant and free radical scavenging activities[5]. Betalains in plants also mitigate damage to photosynthetic capacity in red-pigmented leaves when exposed to excess light, compared to green leaves. Moreover, betalain production is upregulated in response to light or UV radiation[6], and increased betalain accumulation under drought and salt stress conditions protects plants against these environmental stressors, especially in species within the Caryophyllales, which are prevalent in arid and semi-arid environments, as well as saline and alkaline soils[7].

Varies evidences supports prevailing hypothesis that anthocyanins and betalains are mutually exclusive. This includes the wholesale loss, functional loss, and reduced expression of anthocyanin biosynthesis genes, as well as degeneration of their regulatory factors, explaining the absence of anthocyanins in betalain-pigmented species within the Caryophyllales[8]. Furthermore, the evolution of betalain biosynthetic enzymes (e.g., tyrosine hydroxylase and DODA) aligns with the recurrent specialization of these species to betalain synthesis[5, 9]. Betalains are produced in three divergent lineages of organisms: the fungal lineage (e.g., *Amanita muscaria*, *Hygrocybe* and *Hygrophorus* in Basidiomycota)[10]; the flowering plant lineage, particularly in the Caryophyllales[8]; and certain bacterial species (e.g., *Gluconacetobacter diazotrophicus* and the cyanobacterium *Anabaena cylindrica*) [11]. However, no evidence suggests that the Ascomycetes are capable of synthesizing betalains. The secondary metabolites produced by filamentous fungi, particularly ascomycetous species like *Monascus*, *Fusarium*, *Penicillium*, and *Aspergillus*, include a wide range of pigments such as β-carotene, melanins, azaphilones, quinones, flavins, ankaflavin, monascin, anthraquinones, and naphthoquinones, which contribute a broad spectrum of colors, including yellow, orange, red, green, purple, brown, and blue [12]. To date, only one type of violet pigment has been identified in *Aspergillus* species. In *Aspergillus nidulans*, knocking out the SUMO gene triggered the production of the violet anthraquinone asperthecin, which confers resistance to UV radiation [13, 14].

The violet pigment synthesized by *A. sydowii* H-1 has not yet been fully characterized. Previous studies indicated that an insertion mutation in the copper transporter protein prevented the formation of violet pigment. Global untargeted metabolomic analysis identified betanin and cyclo-Dopa 5-O-glucoside in the violet fermentation liquor of *A. sydowii* H-1. In this study, we identified, knocked out, and overexpressed genes involved in the betanin biosynthesis pathway and assessed the activities of tyrosinase (AsTYR), AsDODA and AsLigB. Our findings indicated that the betanin biosynthesis pathway was crucial for colony morphology, conidiogenesis, and the synthesis of violet pigment in *A. sydowii* H-1. Besides, AsDODA and AsLigB catalyzed the formation of 4’5 seco-DOPA from L-DOPA in vitro. This study provides the first description of a potential Ascomycota species capable of producing betalains under controlled culture conditions.

## Materials and methods

### Fungal strains, media, and culture conditions

Strain Activation: *Aspergillus sydowii* H-1, obtained from the Sichuan Province Typical Culture Collection Center, was retrieved from the -80°C refrigerator and inoculated onto PDA (potato 200 g/L, glucose 20 g/L, agar 20 g/L) solid medium. The culture was incubated at 28°C for 3 to 4 days. Fresh spores were scraped off using a cotton swab and suspended in sterile water for further use.

Seed Culture: A 200 μL aliquot of spore suspension was inoculated into 200 mL of seed medium (NaNO₃ 2 g/L, KH₂PO₄ 1 g/L, MgSO₄ 0.5 g/L, KCl 0.5 g/L, FeSO₄ 0.01 g/L, Fructose 30 g/L). The culture was incubated at 28°C with shaking at 180 rpm for 36 hours to obtain fresh seed culture.

Fermentation Culture: A 10 mL aliquot of seed culture was inoculated into 200 mL of fermentation medium (Glucose 5 g/L, KH₂PO₄ 1 g/L, Yeast extract 0.5 g/L, NaCl 0.5 g/L, Peptone 3 g/L, abbreviation as FM). Fermentation was carried out for 8 days at 28°C with shaking at 180 rpm. Three biological replicates were included for each group.

### Bioinformatics analysis

The phylogenetic tree of prephenate dehydratase (PDH), tyrosine aminotransferase (TAT), tyrosinase (TYR), DODA, and LigB was constructed using MEGA X through the maximum likelihood method. The conserved domains of these proteins were predicted using the MEME tool. The tertiary structure of these proteins was predicted with the Swiss-Model tool. Molecular docking between AsTYR and L-tyrosine was performed using AutoDock software. Molecular docking between AsDODA and AsLigB with L-DOPA was performed using AutoDock software.

### Detection of tyrosinase enzyme activity

After four days of cultivation of *A. sydowii* H-1 in liquid fermentation medium, fresh mycelium was harvested, and the fermentation broth was discarded. The mycelium was ground in liquid nitrogen 3-4 times, and the resulting powder was dissolved in 500 μL of tyrosinase extract buffer from the Tyrosinase Activity Assay Kit (Solarbio, BC4055). This solution was used as the sample for assessing intracellular tyrosinase enzyme activity. For extracellular tyrosinase enzyme activity, *A. sydowii* H-1 was cultured on solid fermentation medium for 4 days. Spores were collected using a cotton swab, combined with 500 μL of tyrosinase extract, and the mixture was centrifuged at 8000 rpm for 2 minutes at 4°C. The supernatant was retained as the sample for extracellular enzyme activity assessment.

The crude protein content of the intracellular and extracellular tyrosinase enzyme activity samples was measured using the BCA Protein Concentration Assay Kit (Beyotime, P0012) according to the manufacturer’s instructions. Tyrosinase enzyme activity was determined using the Tyrosinase (Tyr) Activity Assay Kit (BC4055).

### Assessment of violet pigment content

Fermentation broth samples from various strains were collected on days 0, 2, 4, 6, and 8. The fermentation broths were centrifuged at 12,000 rpm for 2 minutes, and the supernatants were analyzed at OD_535nm_ to determine crude violet pigment content.

### Construction of knockout and overexpression strains

Knockouts of AsPDH, AsTAT, AsTYR, AsDODA and AsLigB coding genes were achieved by replacing portions of the target genes with a hygromycin resistance cassette using homologous recombination. The upstream and downstream homologous arms and the hygromycin resistance gene were amplified using primers provided in the supplementary files (Table S3). The three fragments were ligated into the prf-HU2 vector using the 2× Hieff Clone® MultiS Enzyme Premix (Yeasen, 10922ES20). Overexpression strains were constructed by cloning full-length genes (including stop codons) into the prf-HU2-eGFP plasmid. After recombinase ligation, the plasmids were transformed into DH5α, resulting in successful knockout and overexpression vectors.

Both the knockout and overexpression vectors were integrated into the A. sydowii H-1 genome via Agrobacterium tumefaciens-mediated transformation. Vectors were transformed into Agrobacterium AGL-1 competent cell, which was grown to an OD_600_ of 0.8, collected by centrifugation at 5000 rpm, and resuspended in IM medium (KH_2_PO_4_ 1.45 g/L, K_2_HPO_4_ 2.05 g/L, NaCl 0.15 g/L, MgSO_4_·7H_2_O 0.50 g/L, CaCl_2_ 0.067 g/L, FeSO_4_·7H_2_O 0.0025 g/L, (NH_4_)_2_SO_4_ 0.50 g/L, Glucose 2.00 g/L, MES 8.54 g/L, Glycerol 5.00 g/L, 200 μM Acetosyringone (As)) to an OD_600_ of 0.2. The culture was incubated at 28°C with shaking at 180 rpm. Simultaneously, *A. sydowii* H-1 was cultured on PDA medium for 3-4 days. 100 μL of AGL-1 (vector) and 100 μL of spore suspension were mixed and cultured on IM (200 μM AS) solid medium with a 5×5 cm nylon transfer membrane at 25°C in the dark for 2 days. Afterward, the membrane was cut into 0.5×5 cm strips, inverted, and placed on PDA medium (containing 100 μg/mL thaumatin, 100 μM streptomycin, and 200 μM cephalosporin) for 4-6 days at 28°C. To verify the successful knockout and overexpression strains, genomic PCR was performed, and the transformants were rescreened on PDA medium containing 200 μg/mL hygromycin.

### Growth phenotype, Hyphal morphology, and subcellular localization

For growth phenotype and hyphal morphology analysis, 1-3 µL of spore suspension was inoculated at the center of PDA solid medium. A sterile coverslip was placed diagonally into the medium at a 45-degree angle, maintaining a distance of 1.0 to 1.5 cm between the insertion point and inoculation site. The experiment was incubated at 28°C for 4 days. Upon mycelial expansion beyond the coverslip, the slides were removed, and mycelium morphology was examined microscopically.

For subcellular localization, the full-length gene was amplified using the primers listed in Table S3 and inserted into the prf-HU2-eGFP vector. The resulting prf-HU2-gene-eGFP vector was transformed into *A. sydowii* H-1. Transformants were screened and observed using a fluorescence microscope for protein localization.

### RNA extraction and quantitative reverse transcription polymerase chain reaction (qRT-PCR)

Mycelial pellets (5-8) were collected into a sterile 5 mL tube and centrifuged at 12,000 rpm for 2 minutes at 4°C. The pellet was treated with liquid nitrogen for 5 minutes and ground 3-4 times before transferring to a 1.5 mL EP tube. The pellet was resuspended in 1 mL of TRIzol reagent and kept on ice for 5 minutes (do not shake). Then, 200 µL of chloroform was added, followed by incubation at 4°C for 3 minutes and centrifugation at 12,000 rpm for 10 minutes at 4°C. The supernatant (about 500 µL) was mixed with 500 µL of 2-propanol and incubated at 4°C for 10 minutes. The mixture was then centrifuged at 12,000 rpm for 10 minutes at 4°C. The RNA pellet was washed twice with 1 mL of 75% ethanol and centrifuged at 12,000 rpm for 3 minutes at room temperature. After drying for 10 minutes, the pellet was dissolved in 30 µL of diethylpyrocarbonate water. Total RNA extracts were stored at -20°C for up to 1 month.

Reverse transcription was performed according to the Takara PrimeScript™ RT Reagent Kit with gDNA Eraser instructions. qRT-PCR was performed using the Takara TB Green™ Premix Ex Taq™ II and the CFX96 Real-Time PCR Detection System User Guide.

### Tests for stress sensitivity

The response of tyrosine pathway mutants to various stressors was tested using MM (minimal medium) supplemented with 5 mM H₂O₂, 200 µg/mL SDS, 200 µg/mL Congo Red (CR), 1.5 M NaCl, 1.5 M KCl, and 1.2 M sorbitol. These conditions were used to assess the stress tolerance of the mutants.

## Results

### Treatment with tyrosinase inhibitors reduced spore pigmentation and decreased violet pigment production in *Aspergillus sydowii* H-1

Copper plays a pivotal role in the synthesis of fungal conidial pigments, functioning as the active metal center in laccase and copper oxidase enzymes[15]. Preliminary experiments revealed that an insertional mutation in the copper transporter protein AsCptA (the T3 strain) resulted in a white colony phenotype, characterized by abundant aerial hyphae and a complete loss of pigment synthesis (Fig. 1a). This study investigated the effects of gradient concentrations of tyrosinase inhibitors (67.5 µg/mL, 125 µg/mL, 250 µg/mL, 500 µg/mL of kojic acid and tropolone) on A. sydowii H-1. The results indicated both kojic acid and tropolone effectively inhibited conidial pigment deposition, with the highest concentration of tropolone (500 µg/mL) exhibiting a pronounced antifungal effect. However, lower concentrations of the inhibitors did not significantly impact colony growth or morphology (Fig. 1a). The ability to synthesize pigment decreased with increasing concentrations of kojic acid and tropolone, suggesting a potential link between tyrosinase activity and pigment synthesis in *A. sydowii* H-1.

**Fig 1.**
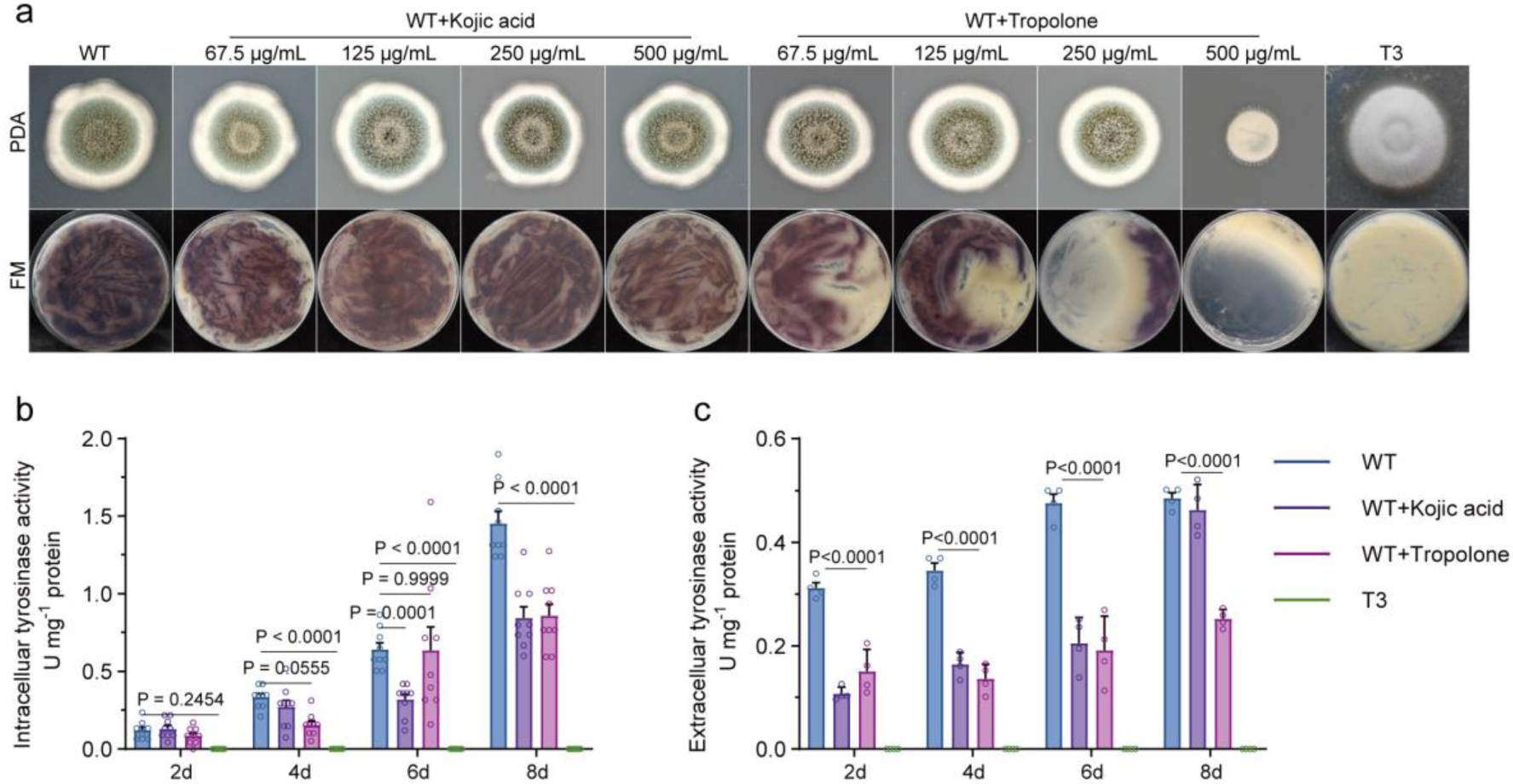
Tyrosinase inhibitors treatment reduced spore pigmentation and decreased violet pigment in *A. sydowii* H-1. (a) Growth and violet pigment phenotypes of *A. sydowii* H-1 exposed to gradient concentrations of tyrosinase inhibitors (67.5 µg/mL, 125 µg/mL, 250 µg/mL, and 500 µg/mL of kojic acid and tropolone). Intracellular tyrosinase activity (b) and extracellular tyrosinase activity (c) in wild type (WT) treated with 500 µg/mL kojic acid and 250 µg/mL tropolone, as well as in the T3 strain.

Fig. 1b illustrated that intracellular tyrosinase activity in *A. sydowii* H-1 increased from 0.07 U/mg crude protein to 1.45 U/mg over 2, 4, 6, and 8 days, whereas extracellular tyrosinase activity rose from 0.31 U/mg crude protein to 0.48 U/mg. Notably, intracellular tyrosinase activity was significantly higher than extracellular activity. Mutations in the copper ion transport protein resulted in a complete loss of tyrosinase activity in the T3 strain. Furthermore, when treated with 500 µg/mL kojic acid and 250 µg/mL tropolone, *A. sydowii* H-1 exhibited significantly reduced intracellular and extracellular tyrosinase activity at all time points (2, 4, 6, and 8 days) compared to the untreated group. These findings indicated a positive correlation between tyrosinase activity and violet pigment synthesis in *A. sydowii* H-1.

### Effects of tyrosine and L-DOPA on spore pigmentation and violet pigment synthesis in *A. sydowii* H-1

To investigate the effects of tyrosine and L-DOPA on spore pigmentation and violet pigment synthesis, *A. sydowii* H-1 was treated with various concentrations of tyrosine and L-DOPA (3.75 mM, 15 mM, 30 mM) (Fig. 2a). DMSO treatment resulted in reduced spore pigment accumulation. Compared to the DMSO group, spore pigmentation decreased with 15 mM and 30 mM tyrosine treatment, but intensified with 15 mM and 30 mM L-DOPA treatment. While DMSO reduced spore pigment deposition, it appeared to stimulate violet pigment synthesis. Tyrosine treatment significantly inhibited pigment synthesis, whereas L-DOPA treatment enhanced it. These results suggested that tyrosinase was involved in the synthesis of both spore and violet pigments. However, the reduction in spore pigment did not significantly affect violet pigment production, implying that spore and violet pigment biosynthesis may proceed through distinct branches of the L-tyrosine metabolic pathway.

**Fig 2.**
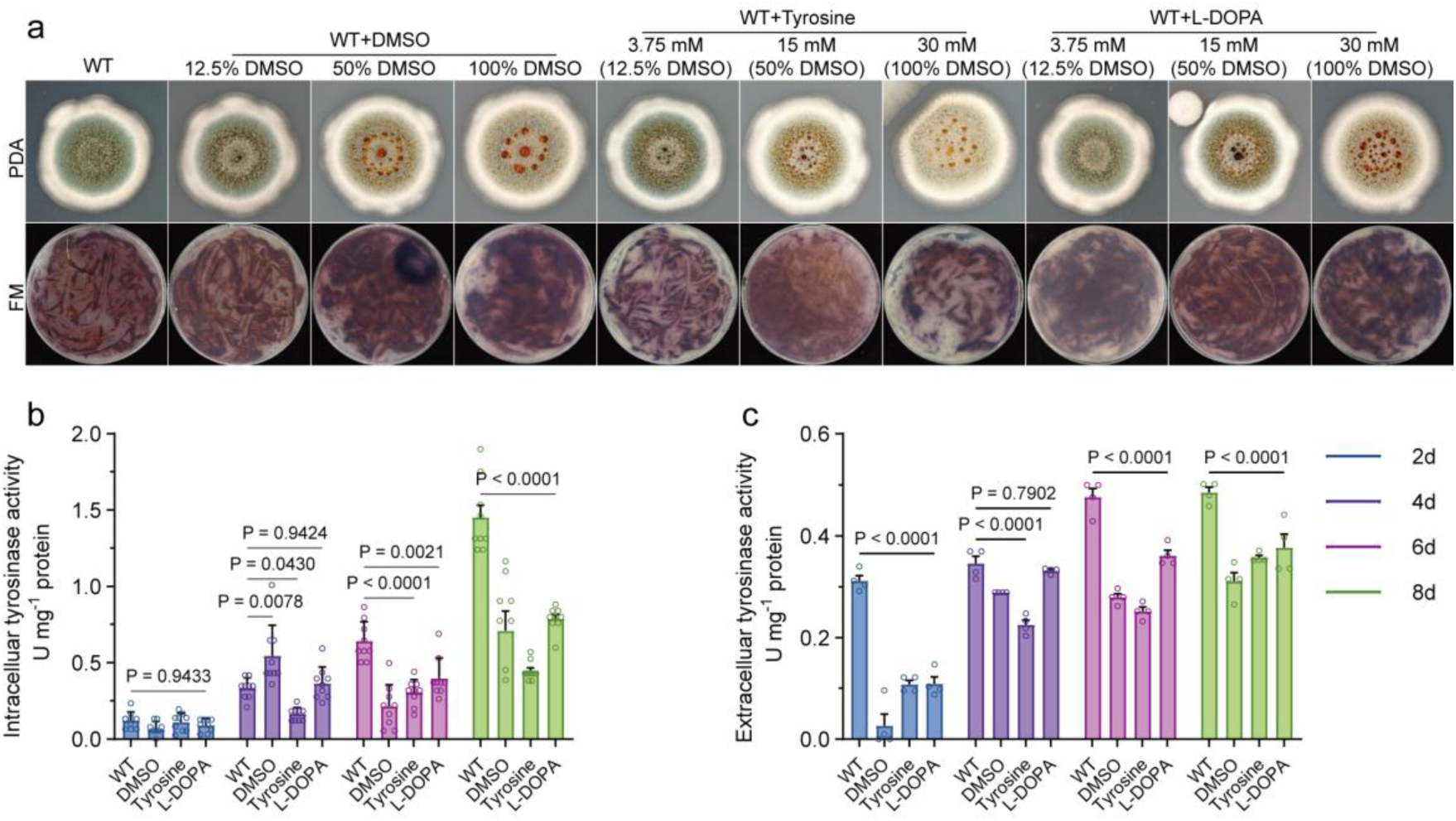
Effects of tyrosine and L-DOPA on spore pigmentation and violet pigment in *A. sydowii* H-1. (a) Growth and violet pigment phenotypes exposed to gradient concentrations of DMSO, tyrosine, and L-DOPA (3.75 mM, 15 mM, and 30 mM). Intracellular tyrosinase activities (b) and extracellular tyrosinase activities (c) in WT treated with 50% DMSO, 15 mM tyrosine and 15 mM L-DOPA.

Next, as shown in Fig. 2b, intracellular tyrosinase activity was measured at 2, 4, 6, and 8 days post-treatment with DMSO. The activity was recorded as 0.06 U, 0.5 U, 0.1 U, and 0.7 U/mg crude protein, respectively, significantly lower than the untreated group. Tyrosine treatment reduced tyrosinase activity, while L-DOPA treatment did not cause any significant changes in enzyme activity. These findings confirmed that tyrosinase activity regulated the growth, development, and secondary metabolism of *A. sydowii* H-1.

### Identification of the Betalain Biosynthesis Pathway in Aspergillus sydowii H-1

Global untargeted metabolomics analysis identified betanin and cyclo-Dopa 5-O-glucoside in *A. sydowii* H-1 violet fermentation liquor (Table S1). To further investigate genes participating in the betanin synthesis pathway, we analyzed the *A. sydowii* H-1 genome and transcriptomic data from fermentation pellets collected at 2 and 8 days (Fig. 5e and Table S2). In the *Aspergillus* genus, tyrosine is not synthesized from L-Arogenate by arogenate dehydrogenase (EC 1.3.1.78); instead, phenylalanine is converted to tyrosine through the catalyzation of prephenate dehydratase (PDH, EC 1.3.1.13) and tyrosine aminotransferase (TAT, EC 2.6.1.5) (As shown in Fig. S10, S11, and S12). Additionally, we identified a CYP450 gene (*EVM0011750.1*) in *A. sydowii* H-1, which had 27.8% sequence similarity with CYP76AD of *Amaranthus tricolor*. However, no other CYP76AD homologs were found in *A. sydowii* H-1, suggesting that tyrosinase (TYR, EC 1.10.3.1) and tyrosine hydroxylase ((EC 1.14.18.1)) may play a role in hydroxylating tyrosine. Further investigation revealed the presence of DOPA 4,5-dioxygenase (DODA, EC 1.13.11-) and two LigB family genes in *A. sydowii* H-1.

We assessed the expression levels of betalain biosynthesis pathway genes in the WT and T3 strain (Fig 3a). Except for the three genes (*EVM0001238.1*, *EVM0003851.1*, and *EVM0002024.1*), the expression levels of other identified genes were markedly evevated compared to the T3 strain. The expression levels of *EVM0011750.1*, *EVM0006581.1*, *EVM0006129.1*, *EVM0004209.1*, *EVM0008360.1*, and *EVM0000393.1* increased progressively with fermentation duration, reaching their peak at 8 days.

**Fig 3.**
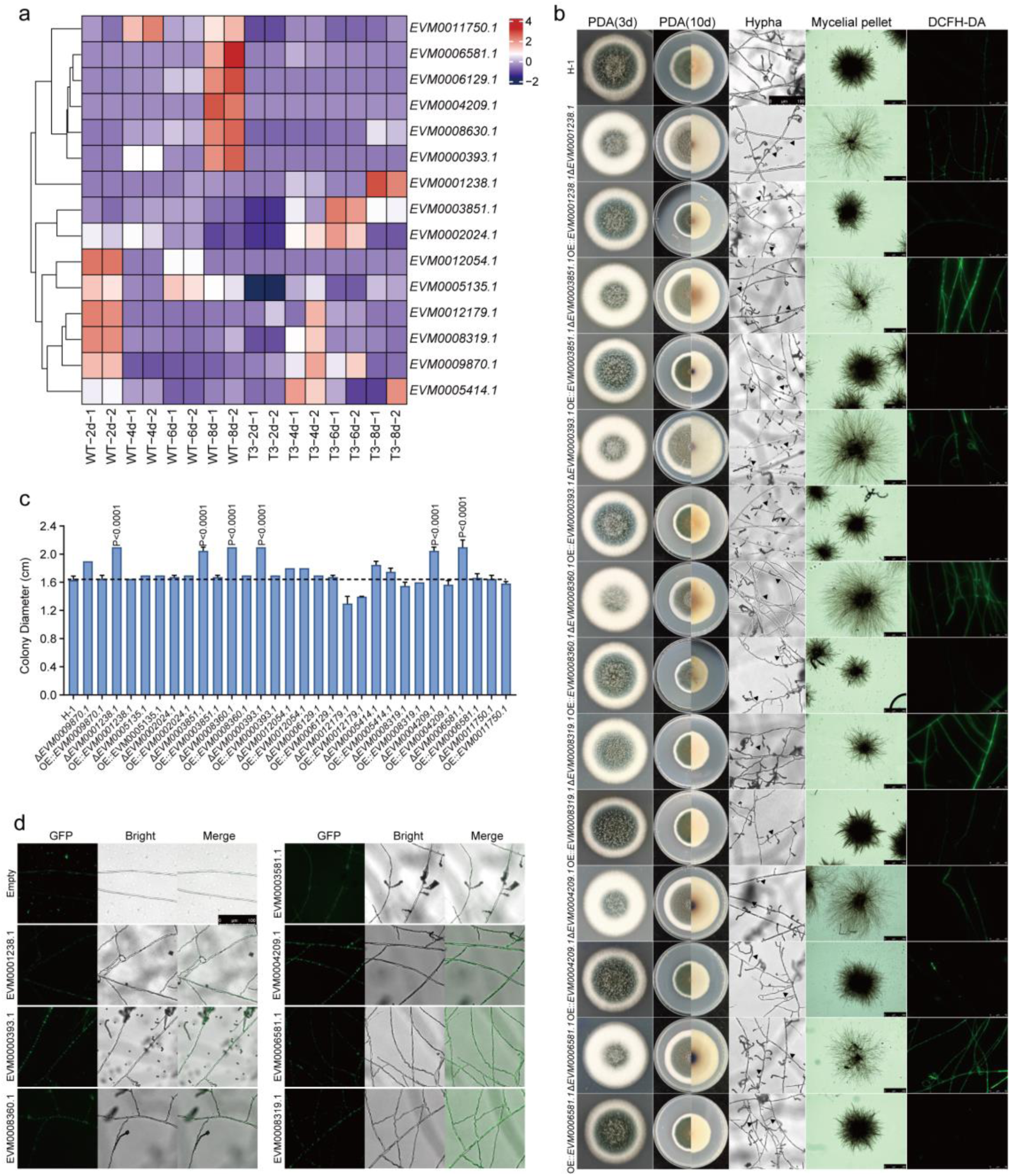
Influence of betalain synthesis pathway genes on the growth and development of *A. sydowii* H-1. (a) Heatmap of the expression levels of betalain biosynthesis pathway genes in the WT and T3 strains. (b) Colony morphology, mycelial pellet appearance, and ROS content of mutants. (c) Colony diameter (cm) of mutants. (d) Subcellular localization of betalain biosynthesis pathway enzymes.

### Influence of betalain synthesis pathway genes on the growth and development of *A. sydowii* H-1

We next constructed knockout and overexpression strains for the genes involved in the betalain biosynthesis pathway. The successful generation of these mutants was confirmed by PCR (Fig. S4 and S5). To assess the impact of these genes on the growth and development of *A. sydowii H-1*, we analyzed the colony morphology and sporulation of the mutantion strains. The T3 knockout strain exhibited reduced conidiation and impaired hyphal development (Fig. S6). Knockout strains for As*TAT* (Δ*EVM0001238.1*), As*TYR* (Δ*EVM0003851.1*, Δ*EVM0000393.1*, Δ*EVM0008360.1*), and As*DODA* (Δ*EVM0006581.1*, Δ*EVM0004209.1*) displayed similar colony morphologies, characterized by dense aerial hyphae, enlarged colony diameters, lighter pigmentation, and notably loose mycelial pellets (Fig. 3b, 3c). In contrast, wild-type (WT) and overexpression (OE) strains formed compact, dark green spores at the colony center, with fewer aerial hyphae at the periphery.

Microscopic analysis revealed an increase in colony diameter and aerial mycelium in the knockout strains (Δ*EVM0001238.1*, Δ*EVM0003851.1*, Δ*EVM0000393.1*, Δ*EVM0008360.1*, Δ*EVM0006581.1*, Δ*EVM0004209.1*), except for Δ*EVM0008319.1*, when compared to the WT and OE strains (Fig. 3b). The conidia of Δ*EVM0001238.1*, Δ*EVM0003851.1*, Δ*EVM0000393.1*, and Δ*EVM0004209.1* strains exhibited abnormal morphology, while the OE strains produced a greater quantity of rounder conidia, with denser, more branched hyphae and increased conidiophore stalk numbers. This suggested that the OE strains may form denser mycelial pellets during liquid fermentation. Additionally, reactive oxygen species (ROS) levels and the GSSG/GSH ratio in the knockout strains were higher than in the WT and OE strains (Fig. S8). Subcellular localization studies indicated that the enzymes encoded by these betalain genes were localized in the cytoplasm and performed their functions there (Fig. 3d). Overall, the genes involved in betalain biosynthesis significantly influenced the colony morphology, growth rate, and sporulation of *A. sydowii* H-1.

### Impact of betalain synthesis pathway genes on the stress response of *A. sydowii* H-1

We subsequently assessed the sensitivity of the mutant strains to various chemical and osmotic stressors in minimal medium (MM) (Fig. 4). Our results showed that exposure to sorbitol stress significantly enhanced the growth of the WT and other mutant strains, although it reduced spore pigment accumulation in the Δ*EVM0001238.1*, Δ*EVM0003851.1*, Δ*EVM0000393.1*, Δ*EVM0008360.1*, Δ*EVM0006581.1*, and Δ*EVM0004209.1* strains. This suggested that genes involved in betalain pigment synthesis were responsive to osmotic stress. Conversely, under salt stress (NaCl and KCl treatments), the growth of the knockout strains (Δ*EVM0001238.1*, Δ*EVM0003851.1*, Δ*EVM0000393.1*, Δ*EVM0008360.1*, Δ*EVM0006581.1*, and Δ*EVM0004209.1*) was significantly impaired compared to that of the WT and OE strains. We also evaluated the response of the mutant strains to oxidative stress (H₂O₂), cell wall stress (Congo red, CR), and cell membrane stress (sodium dodecyl sulfate, SDS). Interestingly, the mutants did not display heightened susceptibility to any of these stress conditions. These treatments appeared to have minimal impact on both the WT and other mutants. Overall, the betalain synthesis pathway genes appeared to significantly influence growth and the response to salt stress, but did not notably affect oxidative, cell wall, or membrane stress responses in *A. sydowii* H-1.

**Fig 4.**
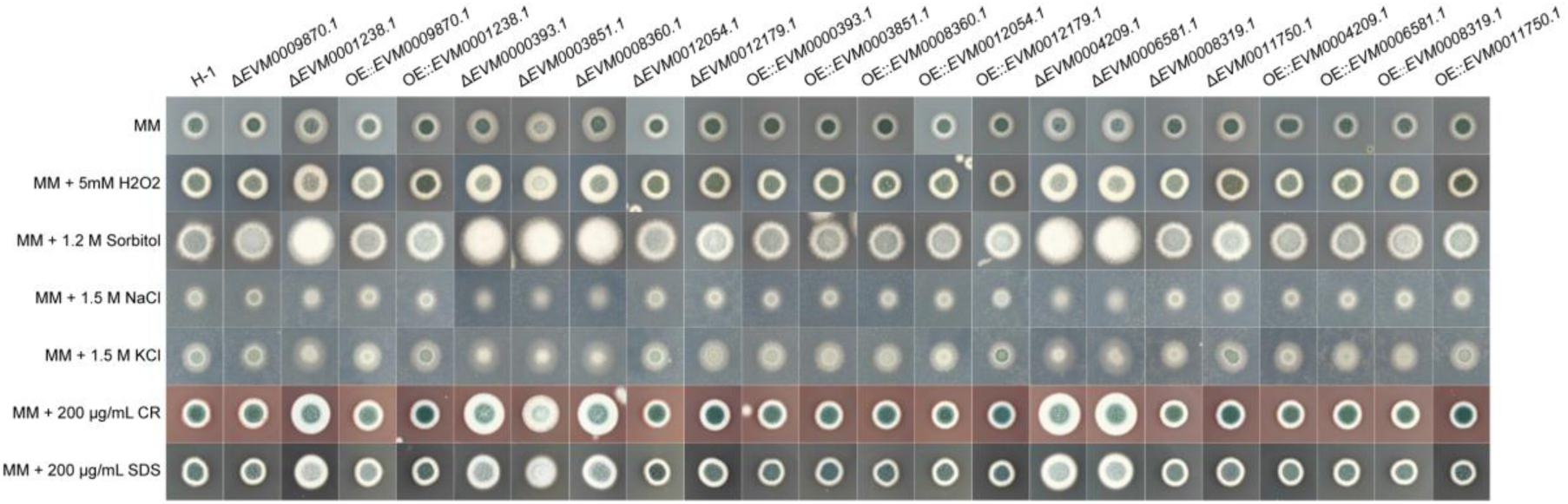
Impact of betalain synthesis pathway genes on the stress response (5 mM H₂O₂, 1.2 M sorbitol, 1.5 M NaCl, 1.5 M KCl, 200 µg/mL CR, and 200 µg/mL SDS) of *A. sydowii* H-1.

### Impact of betalain synthesis pathway genes on violet pigment synthesis in *A. sydowii* H-1

The secondary metabolism of fungi is intricately linked to their growth, development, and sporulation[1]. We evaluated the mutant strains for violet pigment synthesis. Following a three-day cultivation period, the knockout strains of As*PDH*, As*TAT* (except for Δ*EVM0002024.1* and Δ*EVM0005135.1* strains, Fig. S9) and As*TYR* (except for Δ*EVM0006129.1* and Δ*EVM0005414.*1 strains, Fig. S9) could not synthesize pigment, whereas the OE strains showed increased pigment accumulation compared to the WT strain (Fig. 5a, b, and c). Spectrophotometric analysis revealed that the violet pigment exhibited an absorption peak at OD_535nm_. Thus, the pigment content of the fermentation supernatant was measured at λ_535nm_. The maximum pigment synthesis in the WT strain occurred on day eight, reaching an OD_535nm_ of 1.22. The As*PDH* knockout strain (Δ*EVM0009870.1*) produced a brownish-yellow color substance, presumably resulting from flavonoid accumulation. In contrast, the OE::*EVM0009870.1* and OE::*EVM0001238.1* strains accumulated violet pigment more rapidly, reaching an OD_535nm_ of 1.98 and 1.7, respectively, approximately 1.62 and 1.39 times higher than that of the WT strain. Additionally, overexpression As*TAT* led to increasing downstream As*TYRs* expressions (*EVM0003851.1* and *EVM0008360.1*) (Fig. 6b, c, and e). The pigment synthesis capacity of As*TYR* knockout strains exhibited substantial variability. Δ*EVM0003851.1*, Δ*EVM0008360.1*, Δ*EVM0012054.1*, and Δ*EVM0012179.1* strains showed decreased pigment production, with the Δ*EVM0012054.1* fermentation broth exhibiting a yellow coloration. The OD_535nm_ of Δ*EVM0000393.1* strain was 0.24, while Δ*EVM0006129.1* and Δ*EVM0005414.1* strains continued to synthesize pigment (Fig. S9). The OD_535nm_ of OE::*EVM0003851.1*, OE::*EVM0008360.1*, OE::*EVM0012054.1*, and OE::*EVM0000393.1* increased substantially, with values of 3.57, 1.47, 1.64, and 2.74, respectively, corresponding to enhancements of 2.93, 1.20, 1.34, and 2.25 times compared to the WT strain. On days 2, 4, 6, and 8, the As*TYR*s expression levels in the OE::*EVM0000393.1* and OE::*EVM0003851.1* strains rose dramatically, increasing several hundredfold (Fig. 6c, d, e, f, and g). The Δ*EVM0003851.1* and Δ*EVM0000393.1* strains exhibited minimal expression of other As*TYR*s, other than *EVM0012179.1*, perhaps elucidating the absence of pigment production in the absence of As*TYR*s. In summary, overexpression of As*PDH*, As*TAT*, and As*TYR* may raise violet pigment production by increasing metabolic flux in the betalain biosynthetic pathway that contribute to the *A. sydowii* H-1 ecological niche.

**Fig. 5.**
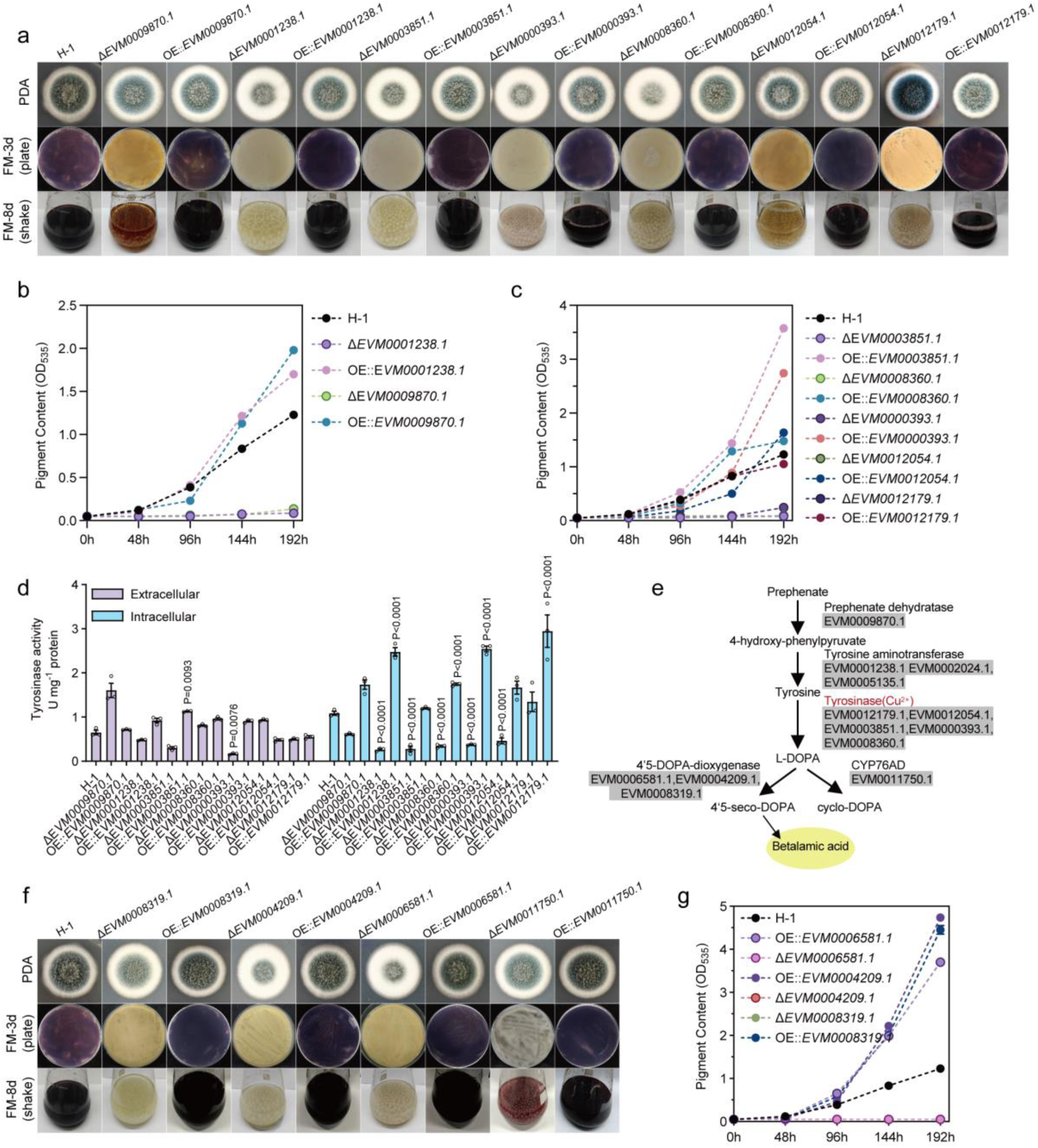
Impact of betalain synthesis pathway genes on violet pigment synthesis in *A. sydowii* H-1. (a) Colony morphology and violet pigment production on solid and liquid fermentation media of As*PDH*, As*TAT*, and As*TYR* mutants. (b) Violet pigment accumulation curves for As*PDH* (*EVM0009870.1*) and As*TAT* (*EVM0001238.1*) deletion and OE strains. (c) Violet pigment accumulation curves for As*TYRs* (*EVM0003851.1*, *EVM0000393.1*, *EVM0008360.1*, *EVM0012054.1*, *EVM0012179.1*) deletion and OE strains. (d) Intracellular and extracellular tyrosinase activities of As*PDH*, As*TAT*, and As*TYR* deletion and OE strains. (e) Hypothetical diagram of the betalain synthesis pathway in *A. sydowii* H-1. (f) Colony morphology and violet pigment production on solid and liquid fermentation media of ilJJ

**Fig 6.**
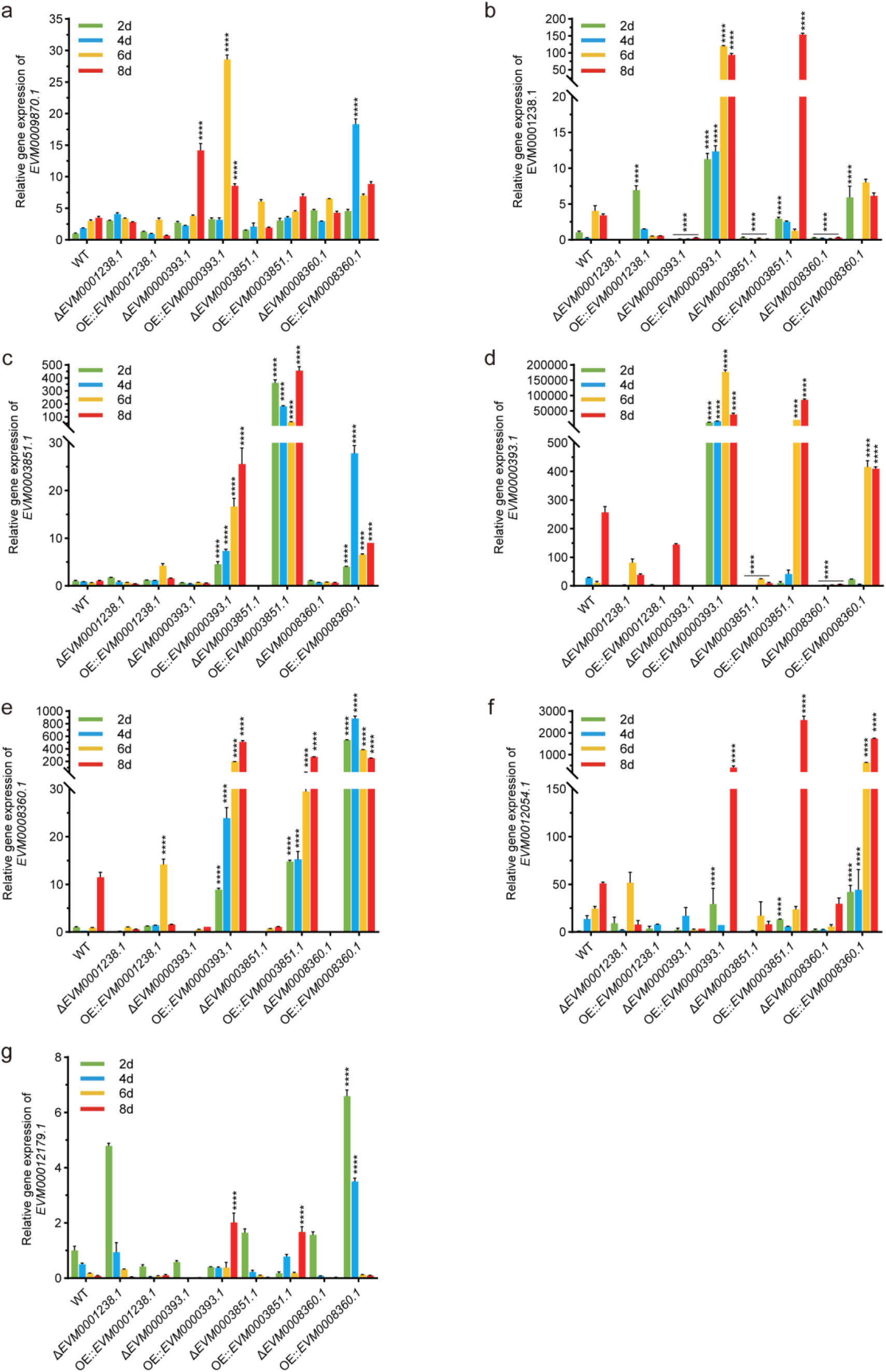
Expression levels of betalain biosynthesis pathway genes in As*TAT* and As*TYR* mutants. (a) Expression level of As*PDH EVM0009870.1*. (b) Expression level of As*TAT EVM0001238.1*. (c) Expression level of As*TYR EVM0003851.1*. (d) Expression level of As*TYR EVM0000393.1*. (e) Expression level of As*TYR EVM0008360.1*. (f) Expression level of As*TYR EVM0012054.1*. (g) Expression level of As*TYR EVM0012179.1*.

We next evaluated the impact of As*PDH*, As*TAT*, and As*TYR* on tyrosinase activity. Except for OE::*EVM0003851.1* and Δ*EVM0000393.1* strains, the extracellular tyrosinase activity of the mutants resembled to the WT strain. Subcellular localization analysis indicated that tyrosinase predominantly functioned in the cytoplasm. Compared to the WT strain, the Δ*EVM0001238.1*, Δ*EVM0003851.1*, Δ*EVM0008360.1*, Δ*EVM0000393.1*, and Δ*EVM0012054.1* strains exhibited intracellular tyrosinase activities of 0.26, 0.28, 0.34, 0.37, and 0.46 times, respectively, while the OE strains showed increases of 2.47, 1.20, 1.75, 2.54, and 1.67 times. These results suggested that tyrosinase activity directly influenceed violet pigment production in *A. sydowii* H-1.

Finally, we investigated the effects of downstream gene alterations in As*TYR* on secondary metabolism. Unexpectedly, deletion of As*DODA* (*EVM0008319.1*) and As*LigB* (*EVM0006581.1* and *EVM0004209.1*) prevented synthesizing pigment, but the OE::*EVM0008319.1*, OE::*EVM0006581.1*, and OE::*EVM0004209.1* strains resulted in OD_535nm_ values of 4.45, 4.74, and 3.70, respectively, which were 3.65, 3.89, and 3.03 times higher than the WT strain. Deletion of *CYP450* (*EVM0011750.1*) did not significantly affect pigment production. We hypothesized that AsLigB increased L-DOPA availability, highlighting its importance in *A. sydowii* H-1 pigment biosynthesis.

### L-DOPA as a key substance of the violet pigment synthesis of *A. sydowii* H-1

To further investigate the role of L-DOPA in the synthesis of violet pigment by *A. sydowii* H-1, we supplemented the culture medium with L-tyrosine and L-DOPA and examined the pigment phenotypes of As*PDH*, As*TAT* and As*TYR* mutations (Fig. 7a and 7b). The results showed that addition of 15 mM L-tyrosine did not restore pigment synthesis in the As*PDH* and As*TAT* deletion strains (Δ*EVM0009870.1*, Δ*EVM0001238.1*, Δ*EVM0005135.1*, and Δ*EVM0002024.1*) (Fig. 7a). However, after adding 15 mM L-DOPA, pigment synthesis was partially restored in some mutants. Specifically, the Δ*EVM0001238.1* and Δ*EVM0008360.1* strains partially restored pigment synthesis after 3 and 5 days of culture, respectively, while the Δ*EVM0003851.1* and Δ*EVM0012054.1* strains showed partial restoration by day 3. Notably, the Δ*EVM0009870.1* and Δ*EVM0000393.1* strains still failed to synthesize pigments, even after L-DOPA supplementation. These findings further confirmed the critical role of L-DOPA in violet pigment synthesis.

**Fig. 7.**
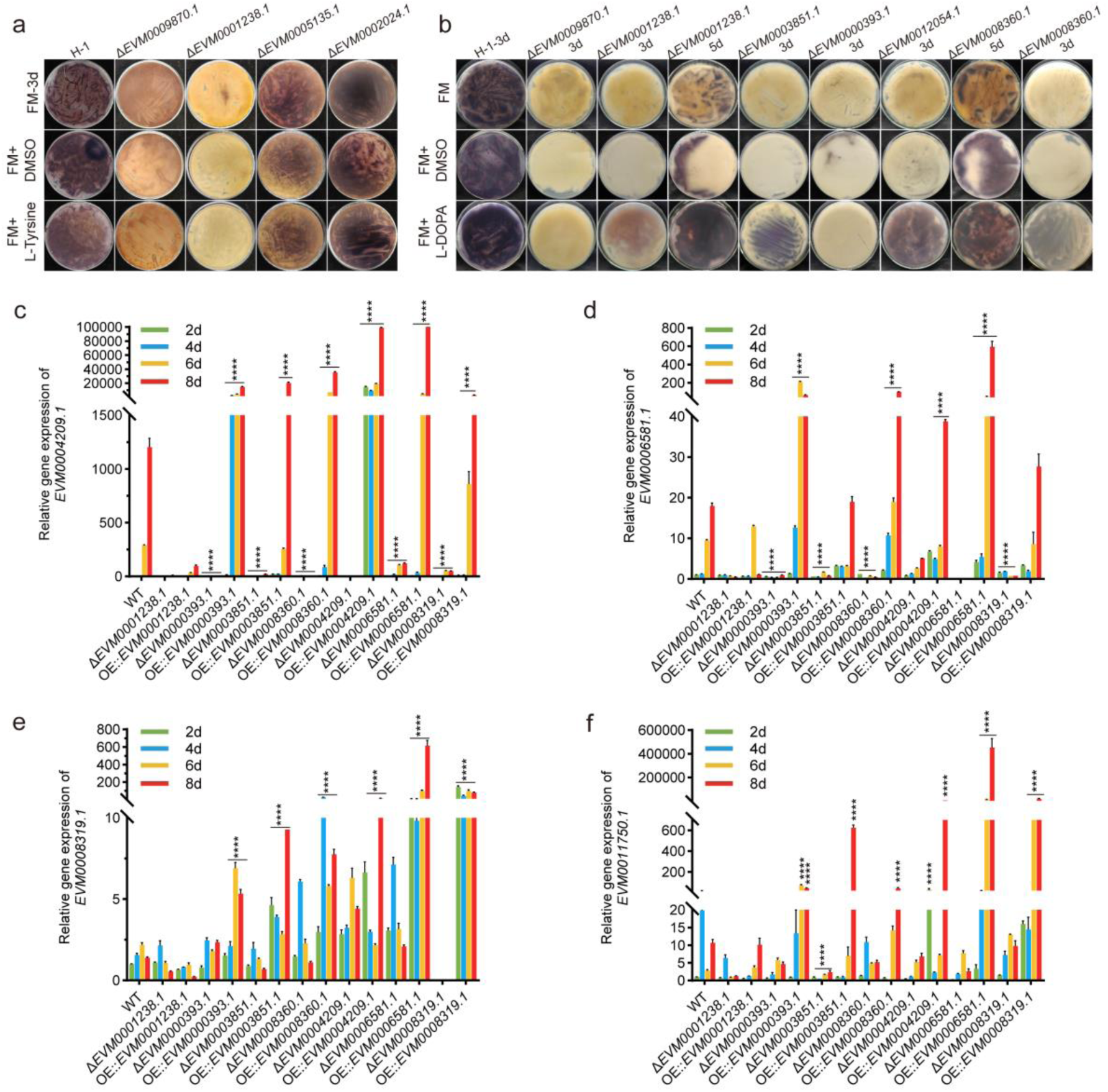
L-DOPA as a key substance in violet pigment synthesis of *A. sydowii* H-1. (a) Violet pigment phenotypes of As*PDH* and As*TAT* deletion strains supplemented with 15 mM L-tyrosine in the culture medium. (b) Violet pigment phenotypes of As*PDH*, As*TAT*, and As*TYRs* deletion strains supplemented with 15 mM L-DOPA in the culture medium. (c) Expression levels of As*LigB* (*EVM0004209.1*) in As*TAT*, As*TYR*, As*DODA*, and As*LigB* mutants. (d) Expression levels of As*LigB* (*EVM0006581.1*) in As*TAT*, As*TYR*, As*DODA*, and As*LigB* mutants. (e) Expression levels of As*DODA* (*EVM0008319.1*) in As*TAT*, As*TYR*, As*DODA*, and As*LigB* mutants. (f) Expression levels of As*CYP450* (*EVM0011581.1*) in As*TAT*, As*TYR*, As*DODA*, and As*LigB* mutants.

We next examined the expression of downstream genes involved in L-DOPA metabolism, including As*DODA*, As*LigB*, and As*CYP450*, in the mutants (Fig. 7c, 7d, 7e and 7f). For the As*LigB* (*EVM0004209.1* and *EVM0006581.1*), the expression levels of these genes increased significantly with the duration of fermentation and pigment accumulation (Fig. 7c and 7d). Compared to day 2, the expression levels of *EVM0004209.1* and *EVM0006581.1* increased by 290-fold and 9.47-fold, respectively, on day 6; the expression levels of these two genes were 1200-fold and 17.9-fold higher, respectively, on day 8. In contrast, in the As*TAT*, As*TYR*, As*DODA* and As*LigB* knockout strains that did not produce pigment, the expression levels of As*LigB* did not significantly increase over time. In the As*TYR*, As*DODA* and As*LigB* OE strains, the expression levels of *EVM0004209.1* and *EVM0006581.1* increased significantly over the fermentation period. On day 6, the *EVM0004209.1* expression level in OE::*EVM0000393.1*, OE::*EVM0008360.1*, OE::*EVM0004209.1*, OE::*EVM0006581.1* and OE::*EVM0008319.1* strains was 14.09, 24.56, 66.26, 15.9, and 2.96 times that of the WT strain, respectively. On day 8, the expression levels of *EVM0004209.1* in OE::*EVM0000393.1*, OE::*EVM0003851.1*, OE::*EVM0008360.1*, OE::*EVM0004209.1*, OE::*EVM0006581.1* and OE::*EVM0008319.1* strains were 12.18, 17.03, 29.37, 82.52, 151.27, and 2.5 times higher than the WT strain, respectively. At the same time, on day 6, the expression levels of *EVM0006581.1* were 21.96, 2.01, and 4.88 times those of the WT in OE::*EVM0000393.1*, OE::*EVM0008360.1*, and OE::*EVM0006581.1* strains, respectively, with day 8 values of OE::*EVM0000393.1*, OE::*EVM0003851.1*, OE::*EVM0008360.1*, OE::*EVM0004209.1*, OE::*EVM0006581.1* and OE::*EVM0008319.1* strains being 3.63, 1.05, 5.56, 2.16, 33.01, and 1.54 times of the WT strain.

For As*DODA EVM0008319.1*, no significant difference in expression levels was observed between the WT strain and the As*TYR* knockout strains (Δ*EVM0000393.1* and Δ*EVM0003851.1*) (Fig. 7e). However, in the As*TYR* OE strains (OE::*EVM0000393.1*, OE::*EVM0008360.1*, and OE::*EVM0003851.1*), the expression level of *EVM0008319.1* was significantly higher than in the WT strain, possibly due to increased intracellular L-DOPA levels. In the As*LigB* deletion strains, *EVM0008319.1* expression level exhibited compensatory upregulation, while in the OE strains (OE::*EVM0004209.1* and OE::*EVM0006581.1*), *EVM0008319.1* expression levels were 2.88 and 45.02 times those of the WT on day 6, and 9.24 and 442.45 times higher on day 8, respectively.

In summary, our results demonstrated that the downstream genes of L-DOPA, including As*DODA* and As*LigB*, exhibitd a marked response during the critical period of pigment synthesis (days 4–8). Overexpression of these genes significantly enhanced the pigment synthesis capacity of *A. sydowii* H-1, further validating the importance of L-DOPA in pigment biosynthesis. Additionally, CYP450 played a key role in the transition of the pigment from colorless to colored. Our data showed that expression of *EVM0011750.1* was significantly higher in the OE strains, particularly in OE::*EVM0004209.1*, OE::*EVM0006581.1* and OE::*EVM0008319.1*.

### EVM0000393.1 and EVM0003851.1 were extremely important tyrosinases of *A. sydowii* H-1

In *A. sydowii* H-1, we identified seven TYRs, and through protein sequence analysis. We confirmed that EVM0000393.1, EVM0003851.1, EVM0008360.1, EVM0012179.1, and EVM0012054.1 all contained Cu(A) and Cu(B) binding sites, essential for tyrosinase activity (Fig. 8a). To determine which of TYRs played a critical role in violet pigment synthesis, we expressed and purified each enzyme in vitro. Activity assays revealed that EVM0000393.1 exhibited the highest tyrosinase activity at 10.55 U/mg protein, with its expression level peaking on day 4 and showing an increasing trend throughout the fermentation process (Fig. 5d). In contrast, the tyrosinase activities of EVM0003851.1 and EVM0012179.1 were 5.74 and 4.99 U/mg protein, respectively, with EVM0012179.1 expression level decreasing as pigment accumulation progressed, suggesting its involvement in the initial stages of fermentation (Fig. 5d). Molecular docking studies further revealed the binding constants of EVM0000393.1, EVM0008360.1, EVM0003851.1, and EVM0012179.1 with L-tyrosine, which were -5.241, -4.667, -4.487, and -4.535 kcal/mol, respectively (Fig. 8c, d, e, f). In vitro assays confirmed that both EVM0000393.1 and EVM0003851.1 exhibited superior tyrosinase activity compared to other enzymes. Furthermore, violet pigment contents were significantly higher in the OE::*EVM0000393.1* and OE::*EVM0003851.1* strains, with 2.25 and 2.93 times greater than those of the WT strain, respectively. These results clearly indicated that EVM0000393.1 and EVM0003851.1 were the most critical tyrosinases involved in violet pigment production in *A. sydowii* H-1.

**Fig. 8.**
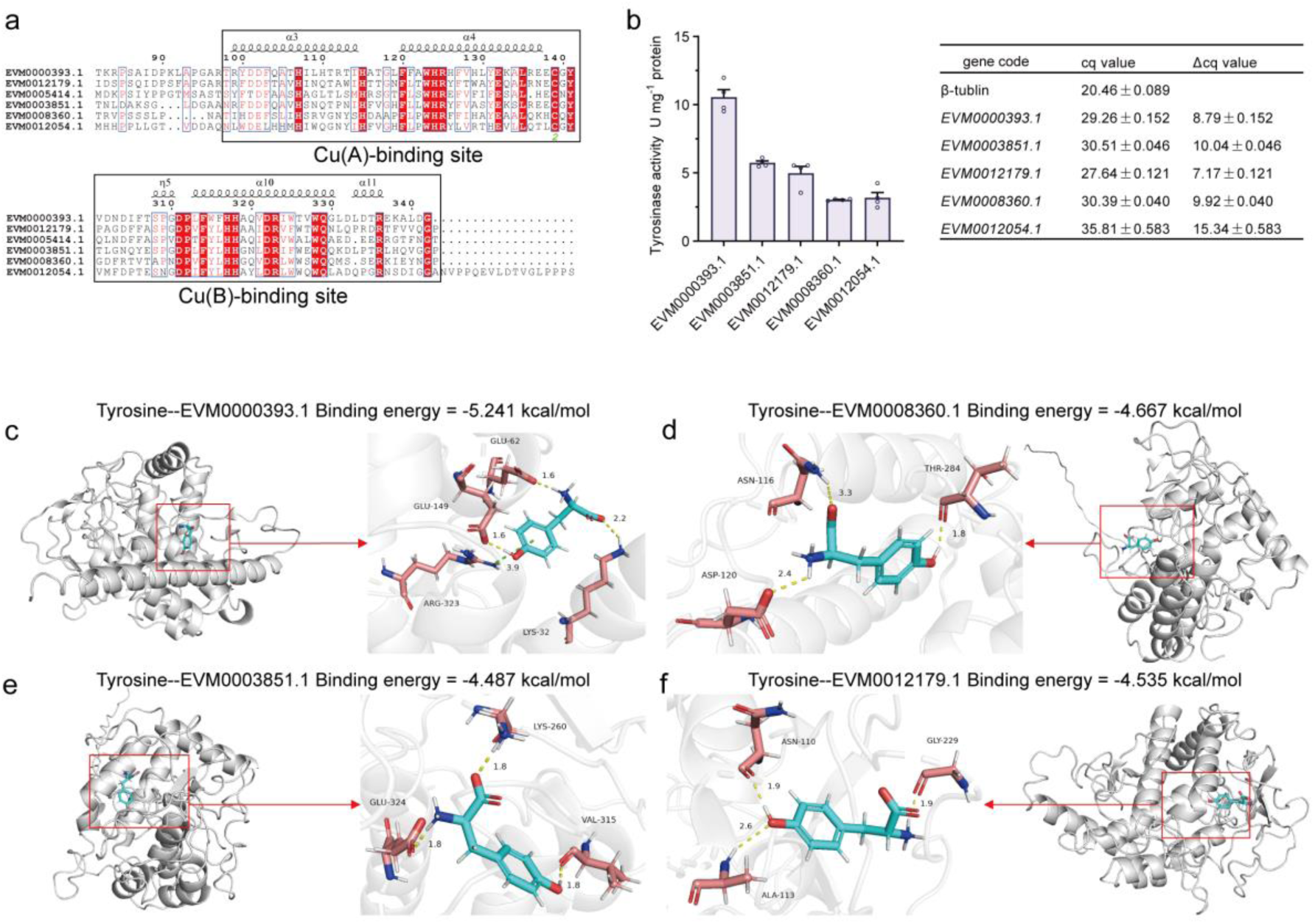
EVM0000393.1 and EVM0003851.1 were crucial tyrosinases in *A. sydowii* H-1. (a) Cu(A) and Cu(B) binding sites of AsTYRs (EVM0003851.1, EVM0000393.1, EVM0008360.1, EVM0012054.1, EVM0012179.1). (b) Tyrosinase activities of expressed and purified AsTYRs in vitro (left) and relative expression levels of As*TYRs* in *A. sydowii* H-1 at day 4 (right). (c) Molecular docking result of EVM0000393.1 and tyrosine. (d) Molecular docking result of EVM0008360.1 and tyrosine. (e) Molecular docking result of EVM0003851.1 and tyrosine. (f) Molecular docking result of EVM0012179.1 and tyrosine.

### Functional characterization of AsDODA and AsLigB

Phylogenetic analysis revealed distinct evolutionary branches for DODA from plants, bacteria, and fungi, with fungal LigB clustering with plant DODA in a single branch (Fig. 9a). The AsLigB proteins (EVM0004209.1 and EVM0006581.1) showed greater similarity to the LigB proteins found in other *Aspergillus* species, while AsDODA (EVM0008319.1) clustered with DODA from Ascomycetes, Basidiomycetes (e.g., *A. muscaria*), and bacteria (e.g., *G. diazotrophicus*). In plants, the *DODA* gene lineage exhibits numerous gene duplication events, the evolutionary implications of which remain ambiguous [9]. Molecular docking analysis showed that the binding constants of EVM0004209.1, EVM0006581.1, and EVM0008319.1 with L-DOPA were -4.170, -4.916, and -3.944 kcal/mol, respectively (Fig. 9b). To confirm their functional roles, we expressed and purified EVM0004209.1, EVM0006581.1, and EVM0008319.1 in vitro. When these purified proteins were added to a reaction mixture containing 10 mM L-DOPA, yellow coloration (λmax 435 nm) was observed exclusively in the AsDODA reaction mixture. High-performance liquid chromatography (HPLC) analysis of the reaction products indicated that EVM0004209.1, EVM0006581.1, and EVM0008319.1 catalyzed the conversion of L-DOPA to the intermediate 4,5-seco-DOPA. Based on these findings, we hypothesized that the yellow pigment may correspond to betalamic acid (Fig. 9c and 9d).

**Fig. 9.**
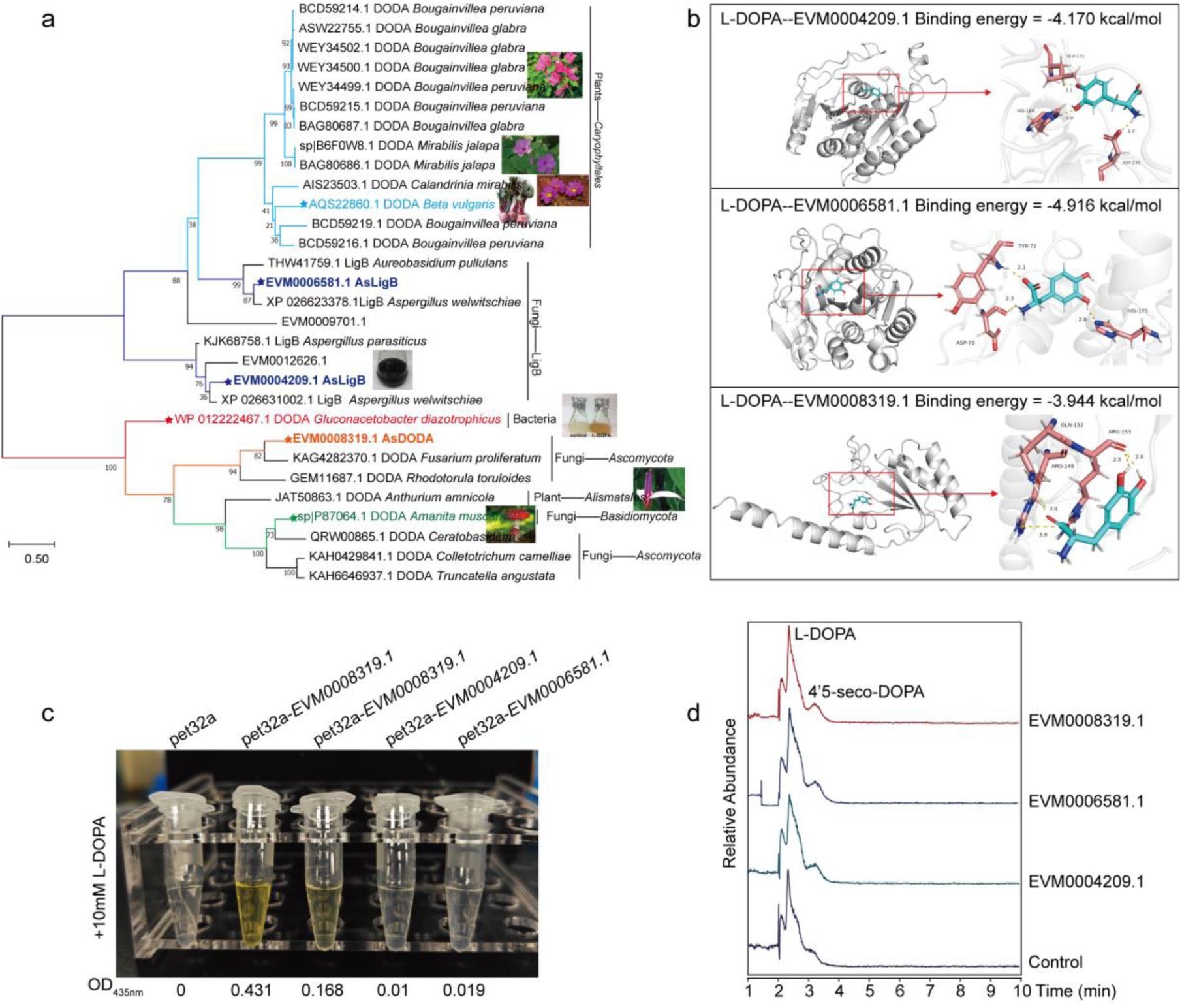
Functional characterization of AsDODA (EVM0008319.1) and AsLigB (EVM0004209.1 and EVM0006581.1). (a) Phylogenetic analysis of DODA and LigB from plants, bacteria, and fungi. (b) Molecular docking results of AsDODA and AsLigB with L-DOPA. (c) Reaction with L-DOPA results of expressed and purified AsDODA and AsLigB in vitro. (d) HPLC analysis of the reaction products of AsDODA and AsLigB catalyzed L-DOPA.

### Transcriptional regulation of betalain genes of *A. sydowii* H-1

In *A. sydowii* H-1, we identified a total of four bHLHs, twelve MYBs, and five WD40s transcription factors. Previous studies in plants have demonstrated that bHLH, MYB, and WD40 transcription factors can assemble into MBW complexes to regulate the synthesis of anthocyanins and flavonoids. In *A. sydowii* H-1, three transcription factors—AsbHLH (EVM0000420.1), AsMYB1R (EVM0011581.1), and AsWD40 (EVM0002833.1)—were found to positively regulate the expression of key betalain biosynthesis genes, including As*TYR* (*EVM0003851.1* and *EVM0000393.1*), As*DODA* (*EVM0008319.1*), and As*LigB* (*EVM0004209.1* and *EVM0006581.1*) (Fig. 10b, c, d, e). These transcription factors also promoted violet pigment production (Fig. 10a). Based on these findings, we proposed that the bHLH, MYB, and WD40 transcription factors in *A. sydowii* H-1 enhanced violet pigment synthesis by upregulating the expression levels of key genes involved in betalain biosynthesis.

**Fig. 10.**
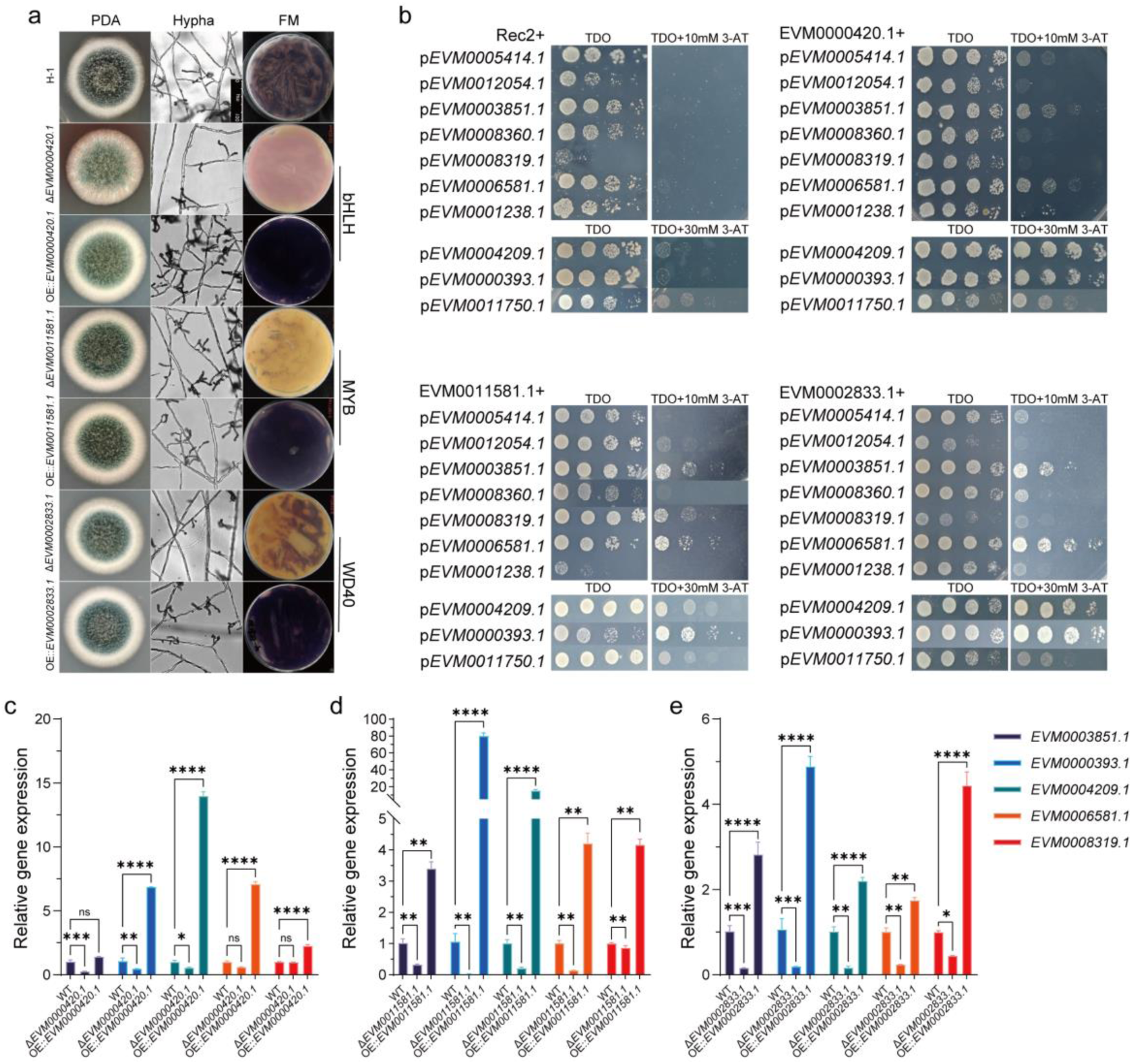
Transcriptional regulation of betalain genes in *A. sydowii* H-1. (a) Colony morphology and violet pigment production on solid and liquid fermentation media of As*bHLH* (*EVM0000420.1*), As*MYB1R* (*EVM0011581.1*), and As*WD40* (*EVM0002833.1*) mutants. (b) Y1H assay of EVM0000420.1, EVM0011581.1, and EVM0002833.1 binding to the promoters of betalain genes. (c) Expression levels of five key betalain genes (*EVM0003851.1*, *EVM0000393.1*, *EVM0004209.1*, *EVM0006581.1*, and *EVM0008319.1*) in As*bHLH* (*EVM0000420.1*) mutants. (d) Expression levels of five key betalain genes in As*MYB1R* (*EVM0011581.1*) mutants. (e) Expression levels of five key betalain genes in As*WD40* (*EVM0002833.1*) mutants.

## Discussion

In *A. sydowii* H-1, we identified and validated a potential complete betalain biosynthesis pathway, which involved several key genes and enzymes, including AsTAT, AsTYR, AsDODA, and AsLigB (Fig. 11). Futhermore, these key genes and violet pigment synthesis were regulated by transcription factors such as AsbHLH, AsMYB1R, and AsWD40.

**Fig. 11.**
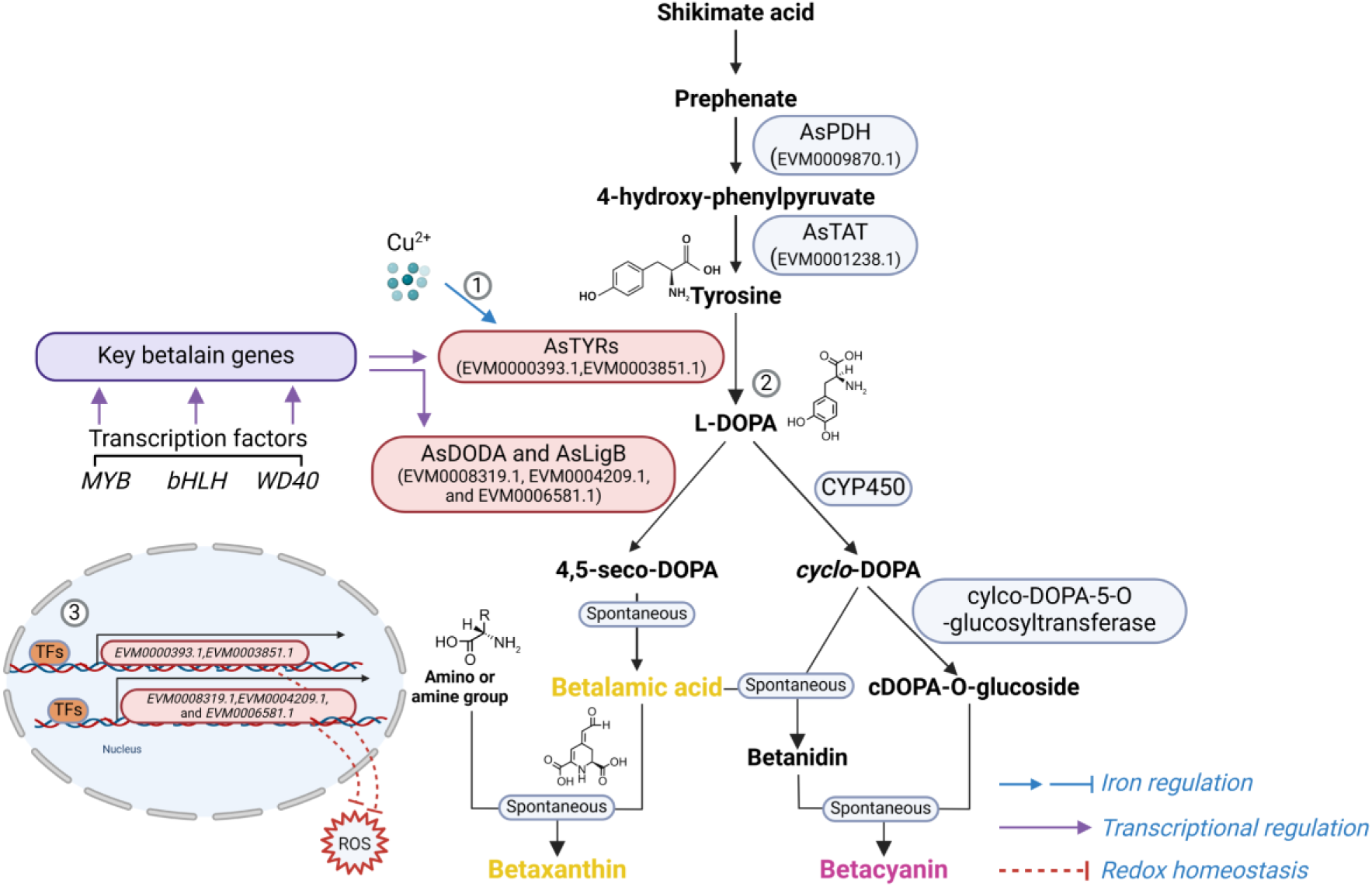
The potential betalain biosynthesis pathway and regulation of betalain genes in *A. sydowii* H-1. ① Cu²⁺, as a static cofactor of AsTYRs, positively influences violet pigment synthesis; ② AsPDH and AsTAT are responsible for converting prephenate to L-tyrosine. AsTYR enzymes (e.g., EVM0000393.1, EVM0003851.1, EVM0008360.1) catalyze the hydroxylation of L-tyrosine to L-DOPA. AsDODA converts L-DOPA to 4’5-seco-DOPA, which may lead to the production of betalamic acid. AsLigB collaborates with AsDODA to catalyze the cyclization of L-DOPA to betalamic acid. ③ AsbHLH, AsMYB1R, and AsWD40 transcription factors regulate the expression of key betalain biosynthesis genes, including AsTYR, AsDODA, and AsLigB. ROS refers to reactive oxygen species. PDH: prephenate dehydratase; TAT: tyrosine aminotransferase; TYR: tyrosinase; DODA: DOPA 4,5-dioxygenase.

To date, no examples of betalain production by Ascomycetes have been reported. This study represented the first description of a potential Ascomycete species capable of producing betalains under controlled culture conditions. Molecular genetics have shed light on the betalain biosynthesis pathway and its evolutionary significance in Caryophyllales. However, our findings suggested that the betalain biosynthesis pathway in *A. sydowii* H-1 differed significantly from that of Caryophyllales. Betanin is synthesized from L-tyrosine via the shikimate pathway. As shown in Fig. S10, S11, S12, S13, S14, and S15, species within the Caryophyllales and *Arabidopsis thaliana* produce tyrosine through arogenate dehydrogenase α (EC 1.3.1.78), whereas *Aspergillus* synthesizes tyrosine through the action of TAT. The next step involves hydroxylation of L-tyrosine to L-DOPA by either a bifunctional CYP450 monooxygenase of the CYP76ADα family or by tyrosine hydroxylase [16]. In *A. sydowii* H-1, we identified one tyrosine hydroxylase (EVM0006129.1), six AsTYRs, and one CYP450 (EVM0011750.1), which shared 27.8% pairwise identity with the CYP76AD of *A. tricolor*. Deletion of AsTAT and AsTYRs led to *A. sydowii* H-1 could not synthesize violet pigment.

DODA, a member of the LigB family, is widespread among land plants and bacteria, despite their divergent sequences and distinct catalytic activity [17]. Many betalain-producing species contain multiple DODA genes. Within Caryophyllales, a gene duplication event in the LigB/DODA lineage led to the formation of the DODAα and DODAβ clades. DODAα catalyzes the L-DOPA ring cleavage, producing 4’5 seco-DOPA, which subsequently forms betalamic acid [18]. In Basidiomycetes (e.g., *A. muscaria*) and bacteria (e.g., *G. diazotrophicus*), DODA catalyzes the conversion of L-DOPA to 2,3- and 4,5-seco-DOPA, which spontaneously cyclize to muscaflavin and betalamic acid [4, 11, 19]. In species that do not produce betalains, DODA may serve other functions. For example, in *Arabidopsis thaliana*, AtLigB catalyzes 2,3-extradiol cleavage of DOPA to synthesize muscaflavin and also converts caffeic acid to iso-arabidopic acid via 2,3-extradiol cleavage[20]. Furthermore, the biosynthesis of arabidopyrones in *A. thaliana* requires AtLigB via a ring-cleavage dioxygenase[21]. Similarly, in *Stizolobium hassjoo*, the phytoalexins stizolobinic and stizolobic acids are synthesized from DOPA via a ring-cleavage dioxygenase [21]. In *A. sydowii* H-1, AsDODA and AsLigB catalyzed the conversion of L-DOPA to 4’5-seco-DOPA in vitro, wheras 2,3-seco-DOPA intermediate was not detected. Additionally, in *A. tricolor*, the expression of AmDODAα1 and AmDODAα2 resulted in a reaction with L-DOPA, producing a bright yellow color indicative of betalamic acid in the reaction mixture [22]; *A. thaliana* containing 35S: AmDOD also produced yellow colouration in flowers and orange red colouration in seedlings when fed L-DOPA [23]. Based on these results, we speculated that the bright yellow substance generated by AsDODA (EVM0008319.1) was likely betalamic acid. Knockout and overexpression of genes involved in the betalain biosynthesis pathway revealed a positive correlation between these genes and violet pigment synthesis. Combined with global untargeted metabolomic analysis and functional characterization of AsDODA and AsLigB, we inferred that *A. sydowii* H-1 had the potential to produce betalains.

Betalains and the more common anthocyanin pigments have never been found together in the same plant[8]. In several plants, the synthesis of anthocyanins is regulated by a highly conserved MYB–bHLH–WD (MBW) transcriptional regulatory complex [24]. Recent studies have revealed that WRKY transcription factors, in conjunction with the MBW complex, also regulate anthocyanin biosynthesis [25]. In betalain-producing plants, several studies have reported that MYB and WRKY transcription factors regulate betalain biosynthesis [26–28]。 For example, in red pulp pitaya, betalain biosynthesis is regulated by the Hu1R-MYB132 transcription factor, which promotes the transcription of Hu*ADH1*, Hu*CYP76AD1-1*, and Hu*DODA1*[29], In *Hylocereus polyrhizus*, the WRKY transcription factor HpWRKY44 also activates the expression of Hp*CytP450-like1*[30]. In Beta vulgaris, BvMYB1 regulates the betalain biosynthesis pathway. However, unlike anthocyanin MYBs, BvMYB1 does not interact with the bHLH members of the anthocyanin MBW complex due to the presence of nonconserved residue [28]. Close biological correlation of pigmentation patterns suggested that betalains might be regulated by a conserved anthocyanin-regulating transcription factor complex--the MBW complex[28]. However, there is little evidence for MBW complex regulating betalain biosynthesis. In *A. sydowii* H-1, we identified three transcription factors—AsbHLH (EVM0000420.1), AsMYB1R (EVM0011581.1), and AsWD40 (EVM0002833.1)—which positively regulated the expression of key betalain biosynthesis genes. However, we have not yet found direct evidence of an MBW complex regulating betalain biosynthesis genes in this species.

Copper (Cu), a static cofactor, is primarily found in oxidoreductases, oxygenases, hydroxylases, and transferases, all of which have flexible active sites designed to optimize electron transfer [15]. We genetically and biologically characterized the AsTYRs. Similar to other TYRs, EVM0000393.1, EVM0008360.1, EVM0003851.1, EVM00012054.1, and EVM00012179.1 each contain Cu(A) and Cu(B) binding sites, along with six conserved histidine residues (Fig. 8a) [31]. The T3 strain (insertional mutation in the copper transporter protein AsCptA) exihibted a white colony and a loss of pigment synthesis phenotype. Based on the importance of TYR in synthesizing melanin and betalain, we speculated that AsTYRs were crucial enzymes for violet pigment synthesis in *A. sydowii* H-1. Our finding indicated that the betanin biosynthesis pathway (including AsTYRs and downstream genes of AsTYRs) was also crucial for colony morphology, conidiogenesis, and the synthesis of violet pigment in *A. sydowii* H-1. Previous studies have identified several genes controlling sporulation and development. For instance, mutants deletion of the *CON1*, *CON2*, and *CON4* genes, which are involved in sporulation, resulted in abnormal conidia morphology and a significant reduction in sporulation rates compared to wild-type strains [32]. Deletion of *mylA* in *A. nidulans* affects stress tolerance, cell wall integrity, and conidial viability [33]. Similarly, in *Magnaporthe oryzae*, the ΔMo*Tyr* mutant showed significantly reduced conidiophore stalk formation, conidia germination, and melanin synthesis, with an impact on both infection and pathogenesis[32]. In *A. sydowii* H-1, Δ*EVM0000393.1*, Δ*EVM0003851.1*, Δ*EVM0008360.1*, Δ*EVM0004209.1*, and Δ*EVM0006581.1* strains exhibited lighter-colored colonies, more developed aerial mycelium, thicker mycelial diameters, reduced conidia numbers, abnormal morphology, and elevated ROS content. In plants, tyrosinase is widely involved in immune responses, abiotic stress tolerance, flavonoid homeostasis, and ROS signaling pathways[34]. Increased expression of polyphenol oxidases (PPOs) in tomato enhances resistance to *Pseudomonas syringae* and *Alternaria solani*, while potatoes show improved resistance to soft rot[35]. Our results also indicated the genes involved in the betalain biosynthesis pathway appeared to significantly influence the response to salt stress in *A. sydowii* H-1.

In fungi, development and secondary metabolism are intricately linked. For example, the novel spore-specific regulator *SscA* controls conidiogenesis in *Aspergillus* species [36] and the KdmB-EcoA-RpdA-SntB chromatin complex coordinates fungal development with mycotoxin synthesis[37]. It is clear that genes involved in fungal development and responses to abiotic and biotic stress are often concurrently regulated with the production of secondary metabolites. Notably, deletion of As*TAT* (Δ*EVM0001238.1*), As*TYR* (Δ*EVM0000393.1*, Δ*EVM0003851.1*, Δ*EVM0008360.1*), and As*LigB* (Δ*EVM0004209.1*, Δ*EVM0006581.1*) not only directly disrupted violet pigment synthesis but also impacted secondary metabolism in *A. sydowii* H-1 by affecting growth and development.

Until now, this study was the first to demonstrate that *A. sydowii* H-1 was a potential ascomycete species capable of producing betalains under controlled culture conditions, and it identified and validated a potential complete betalain biosynthesis pathway, providing preliminary insights into the transcriptional regulation of betalain biosynthesis. Collectively, this research fills a gap in the evolutionary understanding of betalain-producing species. It represents an important step toward developing a viable source of natural betalains, with potential applications in pharmaceuticals, cosmetics, and the food industry.

## Acknowledgments

This work was supported by National Natural Science Foundation of China (32271535, 32071479, 32171473), Natural Science Foundation of Sichuan Province (2024NSFSC0035). We also appreciate Dr.Bo Gao from the Analytical & Testing Center of university for help with LC-MS characterization.

## Data availability statement

All data generated or analyzed during this study are included in this published article and its Supplementary Data.

